# Dynamic Bayesian modeling of the social behavior of *Drosophila melanogaster*

**DOI:** 10.1101/2023.03.02.530782

**Authors:** Kirtan Kalaria, Harshad Mayekar, Dhaval Patel, Subhash Rajpurohit

## Abstract

Organismal behavior has always been a challenge to understand. Insects are one of the amenable systems used to understand behavior. A striking variety of insect behaviors gain support from genetic and physiological studies. *Drosophila*, a widely studied model organism due to its known molecular pathways, has also been popular in behavioral studies. Several behavioral traits in *Drosophila* including mating, locomotion, and oviposition choice have been traced to the neuronal level. Yet, the results of behavioral analyses are equivocal since they often overlook the external milieu, such as social context, which evidentially influences behavior. There have been many attempts to model *Drosophila* behavior, however, all have some fundamental issues like lack of complexity, limitation to isolated organisms, and lack of explainability. Here, we model the behavior of a pair of *Drosophila melanogaster* flies using a novel Dynamic Bayesian Network based approach to better understand behavior in a social context. Two models are proposed, each of which is further used as a predictor for predicting the behavior of both the flies in the pair. They are evaluated on an existing dataset and achieve a remarkable performance: 98.22% and 98.32% accuracy on the two models. Our modeling approach could be applied in predicting animal behaviors in a wide variety of contexts to support existing behavioral studies.

## Introduction

*Drosophila* has been a key model organism for over 100 years to the scientific community. It elicits an unprecedented level of curiosity due to the fact that it is an easy-to-handle, well-established model organism used to seek answers to broader questions ranging from genetics, ecology, evolution, and human biology. *Drosophila melanogaster* is considered an excellent model for studying the primary root of diseases in humans [1]. It is quintessentially employed for a miscellany of genetics based studies on systems biology such as muscle development, immunity, and obesity [2]. Seventy five percent of disease-associated human genes have a functional homolog in *Drosophila melanogaster* [3]. Genes responsible for optical and auditory systems, and neurological disorders in higher order organisms have found comparable genetic wiring in the *Drosophila melanogaster* genome [4]. Of recent, a variety of assays have been performed on *Drosophila melanogaster* to study human diseases. It includes nervous system based studies about perception, learning, and evocation as puzzle pieces to better study human nervous system disorders such as dementia, seizures, and epilepsy [5]. *Drosophila melanogaster* has been a popular choice of a model organism in several similar studies [6–8].

Behavioral neuroscience aims to decode how nervous systems synthesize and combine data from the environment, experience, and internal states [9]. Remarkable advances have been made toward this objective, particularly by leveraging model organisms. They have simpler and well-mapped neural systems with fewer neurons than complex organisms. Studying behavior in isolation would rarely yield interpretations manifesting from environmental interactions. Thus, studying how organisms interact with each other or their abiotic surroundings gives a clearer insight into their behavioral processes. Social behavior is of noteworthy significance as the presence of a social construct stimulates key behaviors associated with cooperation and competition, for example, during mating [10]. Studying the behavior of a large number of flies is unachievable by manual annotation and thus necessitates the utilization of high throughput computing [11]. More recently, advanced computer vision tools and the ability to process large datasets have made quantitative studies feasible [12].

Recent advances include unsupervised classification of behavior followed by manual annotation of the obtained clusters. Though such methods for studying animal behavior are relatively new, the results are robust [13]. In the case of *Drosophila melanogaster*, the approach came to fruition when a research identified 122 behavioral states using unsupervised classification [14]. Their methodology was subsequently utilized in another study to capture the behavior of paired flies. This was achieved by tracking the flies individually using computer vision, segmenting the video, and classifying the behavior. Additional information such as distance from each other, and angle of orientation was also captured. Their work on behavioral analytics identified the effect of the distance between the flies, the relative orientation of their partners, and their pair type on behavior [15].

However, little has been done towards modeling the behavior. The existing work on modeling uses Markov chains and ethograms to model behavioral transitions [15, 16]. However, it is too simplistic and thus fails to capture the complex patterns and relationships between variables. Our work aims to advance the efforts by modeling the behavior of paired *Drosophila melanogaster* on the spatiotemporal scale incorporating social and environmental parameters. We propose a trio of probe-mirror-combine method along with Dynamic Bayesian networks, which can be used by biologists as a general modeling tool for various scenarios. The network is used as a predictor of the behavior of both flies and is evaluated.

## Dataset

We used a publicly available dataset [17] from a previous work [15] to study the social behavior of *Drosophila melanogaster* in a paired social context. The data is temporal and spans all three possible contexts - male pair, female pair, and courtship pair (a male and a female). For each pair type, they conducted a number of experiments, as described in Table 1. In each experiment, they released a pair of flies into a circular arena (radius *R* = 11mm) covered with a transparent dome and recorded videos.

**Table 1.**
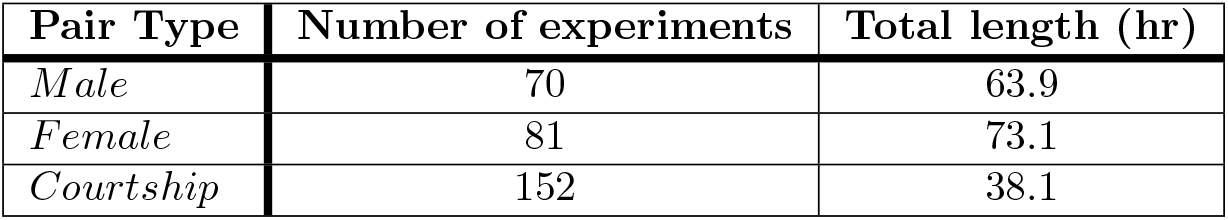
Number of experiments and total video length for each pair type.

| Pair Type | Number of experiments | Total length (hr) |
| --- | --- | --- |
| *Male* | 70 | 63.9 |
| *Female* | 81 | 73.1 |
| *Courtship* | 152 | 38.1 |

For each experiment, they applied unsupervised clustering based behavioral quantification to the recorded video. This yielded the following information for each fly in the pair at periodic time intervals:

1. **Body centroid coordinates**: The x and y coordinates are given for the centroid of the body of each fly. The coordinates are given with the center of the arena as the origin.
2. **Heading**: The heading is defined as the angular deviation in either direction from the perfect alignment, i.e., facing the partner fly directly. In other words, it is the angle subtended by the axis of the fly on the line connecting the pair.
3. **Behavior** The detected behavior via the unsupervised approach is refined down to 8 defined coarse behaviors. The behaviors considered and their integer codes in the dataset are displayed in Table 2.

**Table 2.**
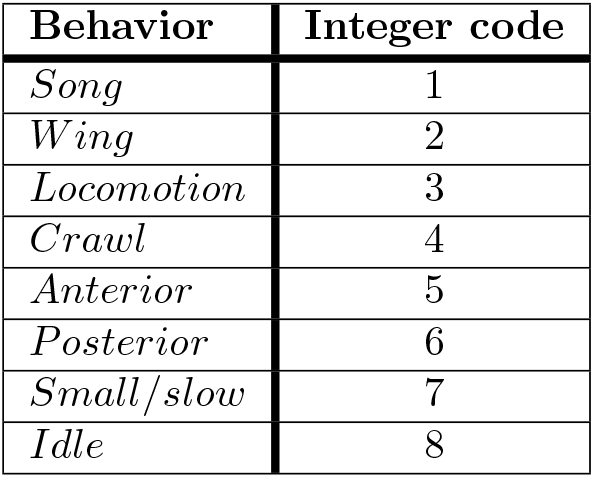
The behaviors considered and the integer encoding in the dataset.

## Preprocessing and Feature Extraction

### Initial Preprocessing

Intensive preprocessing was performed on the dataset to tailor it to the needs of our tasks. For both datasets, the following initial preprocessing steps were carried out:

1. The information of the pair type was included as the sex of each fly to generalize the modeling so that the data from different social contexts could be merged. This allows modeling the effect of the pair type or the partner fly’s sex on the fly behavior.
2. From the location of the flies, meaningful variables were calculated to model the behavior meaningfully. The coordinates data was replaced with the distance between the two flies in a pair, *d_p_*, and the distance of the flies from the center of the arena, *d*_*c*1_ and *d*_*c*2_. Fig 1 illustrates the distances and the headings with respect to the experimental setup.
3. We would like to be able to label either of the two flies as ‘Fly 1’ and the other as ‘Fly 2’. Therefore, for each row in the given dataset, a new row is added to it with the corresponding variables of ‘Fly 1’ and ‘Fly 2’ swapped to reflect the choice of assignment arbitrary in the dataset.
4. All continuous variables were discretized by binning and assigning each bin with an integer code.

**Fig 1.**
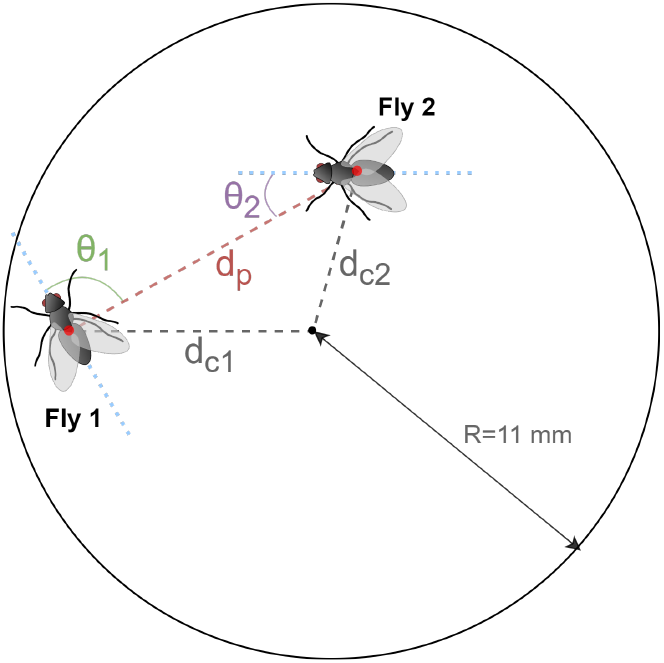
Illustration of the distances *d_p_, d*_*c*1_, and *d*_*c*2_, and the headings *θ*_1_ and *θ*_2_ in the experimental setup. The fly pair is shown in the circular arena. The measurements for the headings and distances are along with the associated variables.

To explore the Bayesian Network structure with a focus on the behavior of one fly, a ‘probe dataset’ was constructed. The Bayesian Network was then manipulated to obtain a faithful representation of the behavior of both the flies in the pair. To fit the final Bayesian Network, a ‘learning dataset’ was constructed that included two-time-step temporal information.

### Preparing the probe dataset

The purpose of the probe dataset is to estimate an understanding of the Bayesian Network structure describing the effect of the environment, including the presence of the partner fly, on the behavior of a fly. Arbitrarily, ‘Fly 1’ was chosen as the fly in focus. The distance from the center (*d_ci_*), the distance between the pair (*d_p_*), the heading from the partner(*θ_i_*), and the sex of the partner (*s_i_*) have shown to be significantly correlated with the behavior of the flies [14].

The distance of the partner fly ‘Fly 2’ from the center of the arena (*d*_*c*2_) defines the partner fly’s physical environmental context and thus has no bearing on the behavior of ‘Fly 1’. Hence, the column corresponding to *d*_*c*2_ was dropped while constructing the probe dataset.

The behaviors of the two flies are correlated and affect one another [14]. However, considering our aim to model the behavior of a pair of flies in one Bayesian Network and keeping in line with the liberty of arbitrary assignment of fly numbers to the flies, the obtained Bayesian Network must have a reflective symmetric structure. Failing to satisfy this condition would result in different representations on the two different assignments. With this constraint, if there exists a directed path from the behavior of ‘Fly 1’, *B*_1_, to the behavior of ‘Fly 2’, *B*_2_, the alternative assignment would demand a reversed directed path, i.e., a directed path from *B*_2_ to *B*_1_. The existence of both such paths would produce a cyclic path, violating the requirement of acyclicity of Bayesian Networks. This argument is also valid for the case where a direct edge exists between the two nodes, instead of a directed path.

Formally, if the directed path from *B*_1_ to *B*_2_ is formed by the set of nodes 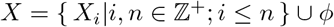 then there will be a directed path from *B*_2_ to *B*_1_ consisting of the set of nodes 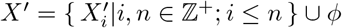 where *X_i_* and 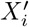 are reflections of each other (the same variable for different flies) in the symmetric network. Fig 2 illustrates the symmetry versus acyclicity dilemma.

**Fig 2.**
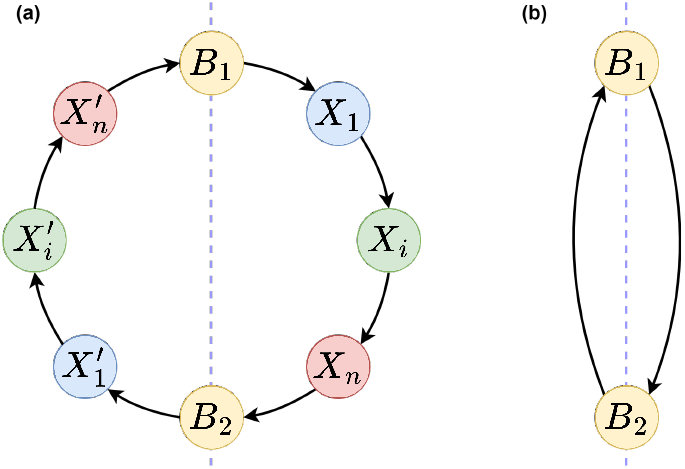
Illustration of the symmetry versus acyclicity dilemma. The cyclic paths formed in the case of symmetric networks are shown for (a) the case of a directed path with intermediate nodes, and (b) the case of a direct edge.

Such a relationship between two modes can be termed a feedback loop. We resolved this dilemma using Dynamic Bayesian Networks. Further, we did not incorporate the effect of *B*_2_ on *B*_1_ while probing, and hence the column corresponding to *B*_2_ was dropped from the dataset. This is rectified later.

In summary, we obtained the probe dataset after dropping *d*_*c*2_ and *B*_2_ in addition to the initial preprocessing steps specified above. The variables constituting the probe dataset are displayed in Table 3. The probe dataset consists of about 54.84 million rows.

**Table 3.** Variables in the probe dataset.

| Variable | Description |
| --- | --- |
| Variable | Description |
| $s_1$ | Sex of ‘Fly 1’ |
| $s_2$ | Sex of ‘Fly 2’ |
| $d_p$ | Distance between ‘Fly 1’ and ‘Fly 2’ |
| $d_{c1}$ | Distance of ‘Fly 1’ from the center of the arena |
| $d_{c2}$ | Distance of ‘Fly 2’ from the center of the arena |
| $\theta_1$ | Heading of ‘Fly 1’ |
| $\theta_2$ | Heading of ‘Fly 2’ |
| $B_1$ | Behavior of ‘Fly 1’ |
| $B_2$ | Behavior of ‘Fly 2’ |

### Preparing the learning dataset

The purpose of the learning dataset is to learn the parameters of the final Dynamic Bayesian Network describing the paired *Drosophila melanogaster* behavior domain.

For this, in addition to the initial preprocessing steps described above, *d_ci_, θ_i_, B_i_*, and *d_p_* from the previous time step are included. They were named 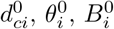, and 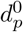 and the current time step *d_ci_, θ_i_, B_i_*, and *d_p_* are renamed 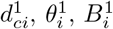, and 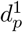. This allows us to use a Dynamic Bayesian Network as it models variables over a temporal scale. The learning dataset also consists of about 54.84 million rows. Fig 3 explains the overall flow of preprocessing to create the two datasets.

**Fig 3.**
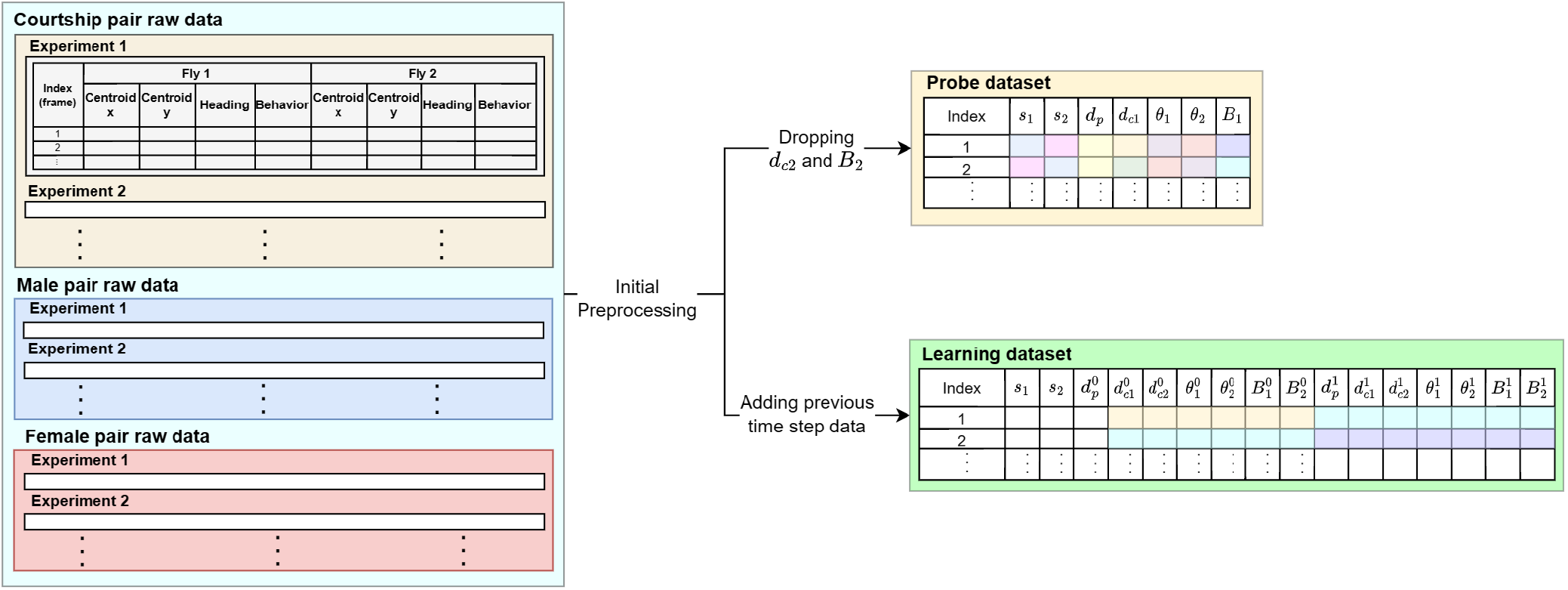
Preprocessing pipeline. The raw dataset is shown on the left from which the probe and learning datasets are formed. The probe dataset is color coded to show double entries of each row in the raw dataset with choice of each fly in the pair as ‘Fly 1’ to make the representation arbitrary in the model. The learning dataset is also color coded to show the formation of temporal features with consecutive observations from the raw dataset.

## Modeling with Bayesian Networks

The proposed prediction model was carefully designed as a representation of the dataset with domain-relevant logical constraints.

Firstly, the structure of the Bayesian network representing the effect of the variables on the behavior of one fly, 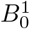, was probed, or learned by applying PC, a constraint-based structure learning algorithm, on the probe dataset. The probed Bayesian Network gives us the domain-relevant structured dependencies between the variables in the form of edges between the nodes, as learned from the data. The PC algorithm builds the structure based on conditional independence tests, taking into account the following prior specified independencies:

- *s*_1_ ⊥ *s*_2_: The sex of ‘Fly 1’ is independent of the sex of ‘Fly 2’. It is an experimental choice of pair type, hence the two variables must be independent of each other.
- *d_p_* ⊥ *d*_*c*1_: The distance between the pair of flies is independent of the distance of each fly from the center of the arena because, with the same distance from the center, the distance between the pair can vary in an uncorrelated manner.

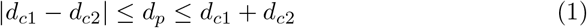

The network formed was termed as the ‘probe network’, denoted by 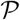. Since the choice of the assignment of the fly number to both flies was arbitrary, all variables of Fly 1 can be swapped with those of Fly 2 and vice versa in the network. *s*_1_, *s*_2_, *d*_*c*1_, *θ*_1_, *θ*_2_, and *B*_1_ were replaced with *s*_2_, *s*_1_, *d*_*c*2_, *θ*_2_, *θ*_1_, and *B*_2_ respectively to yield a new Bayesian network, termed as the ‘mirrored network’ and denoted by 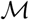. It is as valid a representation of the domain data as the probed network.

In order to obtain a combined representation of the relationships between variables of both flies, we merged the probed network and the mirrored network by including all nodes and edges from both networks into one network termed as the ‘combined network’ and denoted by 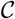. If 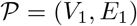 and 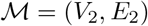:

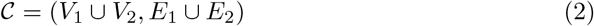

Probing, mirroring, and combining method trio enabled us to represent the paired fly interactions using a symmetric Bayesian network. Fig 4 illustrates the same.

**Fig 4.**
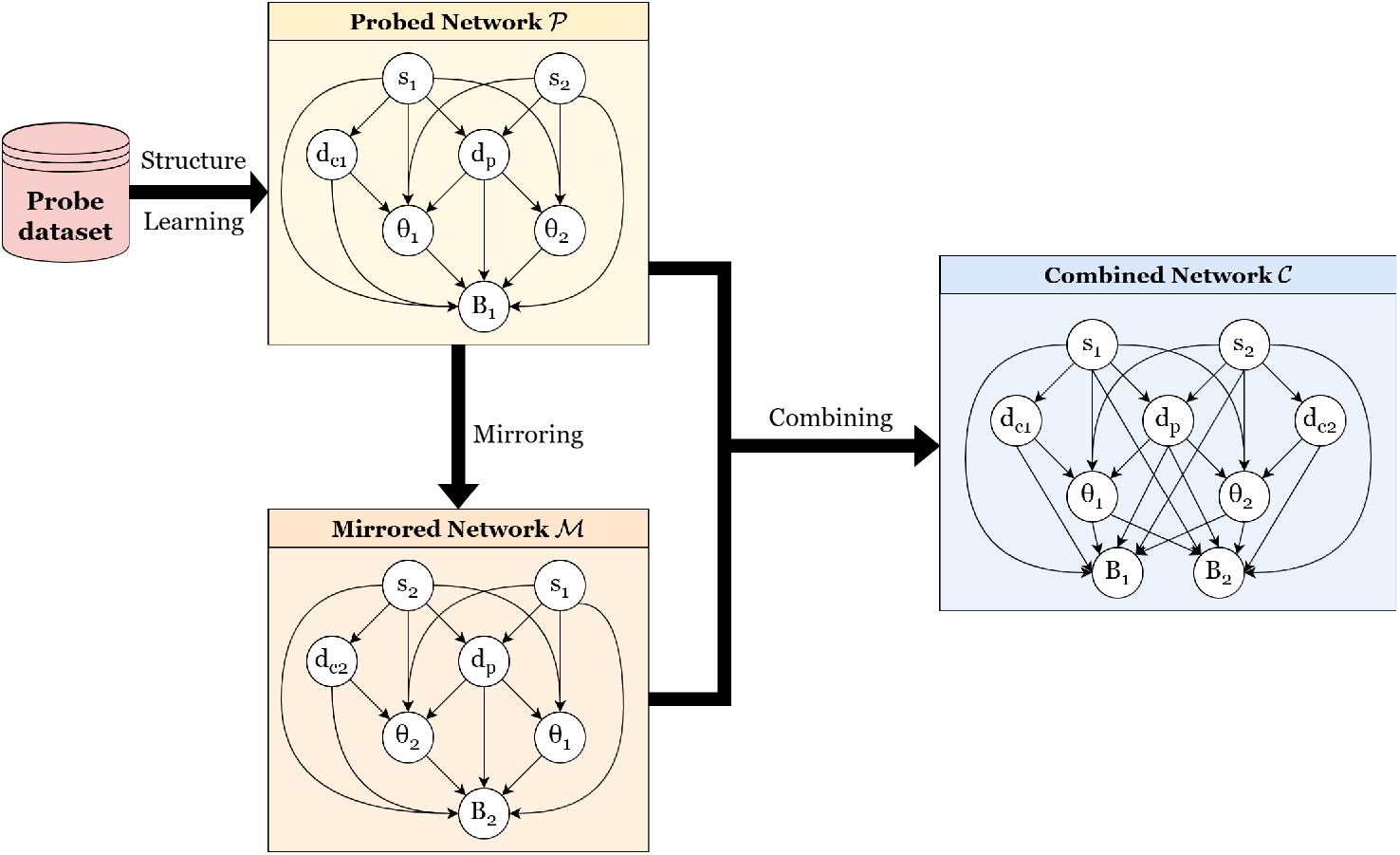
Probing, mirroring, and combining methods. The trio of probing, mirroring, and merging methods is exhibited on our model.

The method that allows us to prevent the violation of acyclicity is the use of Dynamic Bayesian Networks. They overcome the shortcoming of the regular Bayesian networks to model the cyclic dependencies found in nature by enabling connections in the temporal domain, i.e., across time slices and not within the same time slice. Since the behavior of one fly affects that of the other, connections from *B*_0_ to *B*_1_ and from *B*_1_ to *B*_0_ were established in the temporal domain. Dynamic Bayesian Networks (DBNs) are a subclass of Bayesian Networks that include variables over multiple time steps. They help find the relations of the variables on the variables from adjacent time steps. They also uphold the Markovian assumption and generalize Hidden Markov Models (HMMs). Fig 5 shows a general example of a DBN.

**Fig 5.**
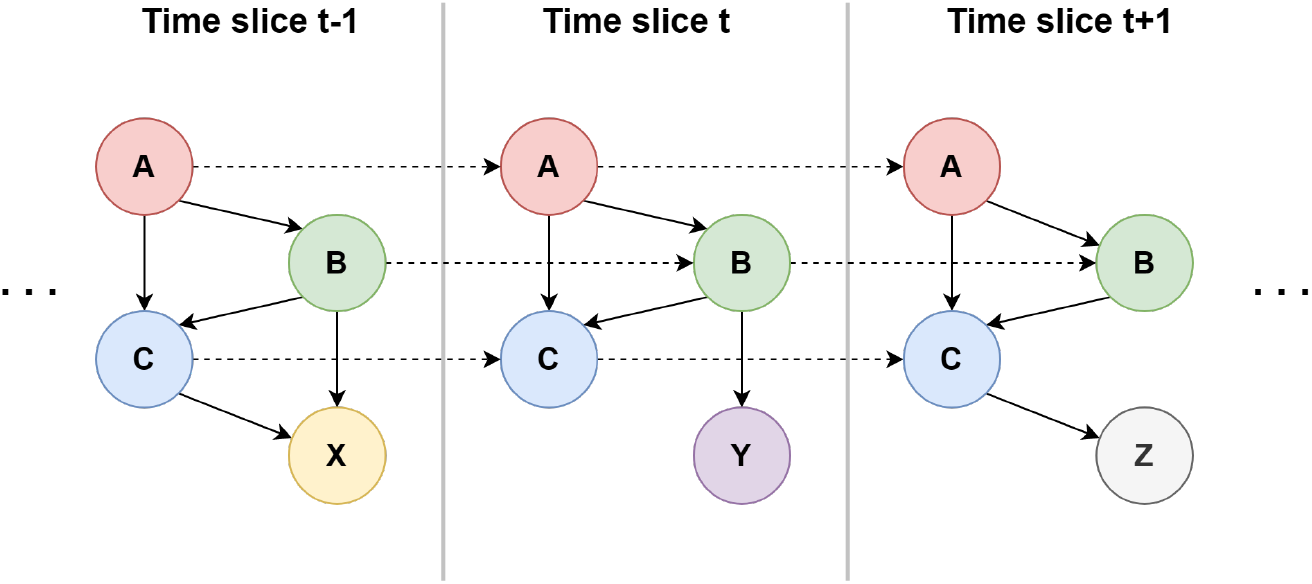
Dynamic Bayesian Networks. Dynamic Bayesian Networks are generalized models that have multiple time slices and the edges are allowed within the same time slice as well as across time slices.

We propose two models – the ‘base network’ and the ‘pruned network’ each of which is a 2-Time-Slice Bayesian Network (2TBN) employed to model temporal independencies. For the base network, each time slice consists of the combined network 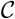. Since the behavior of one fly affects that of the other, connections from *B*_0_ to *B*_1_ and from *B*_1_ to *B*_0_ were established in the temporal domain, and the Markovian connections were upheld for all variables. The sex of the flies *s*_0_ and *s*_1_ were omitted from the first time slice as they will remain constant and are unchanging parameters. Thus, DBNs were leveraged to overcome our acyclicity versus symmetry dilemma, as a final method of constructing our proposed network, as shown in Fig 6.

**Fig 6.**
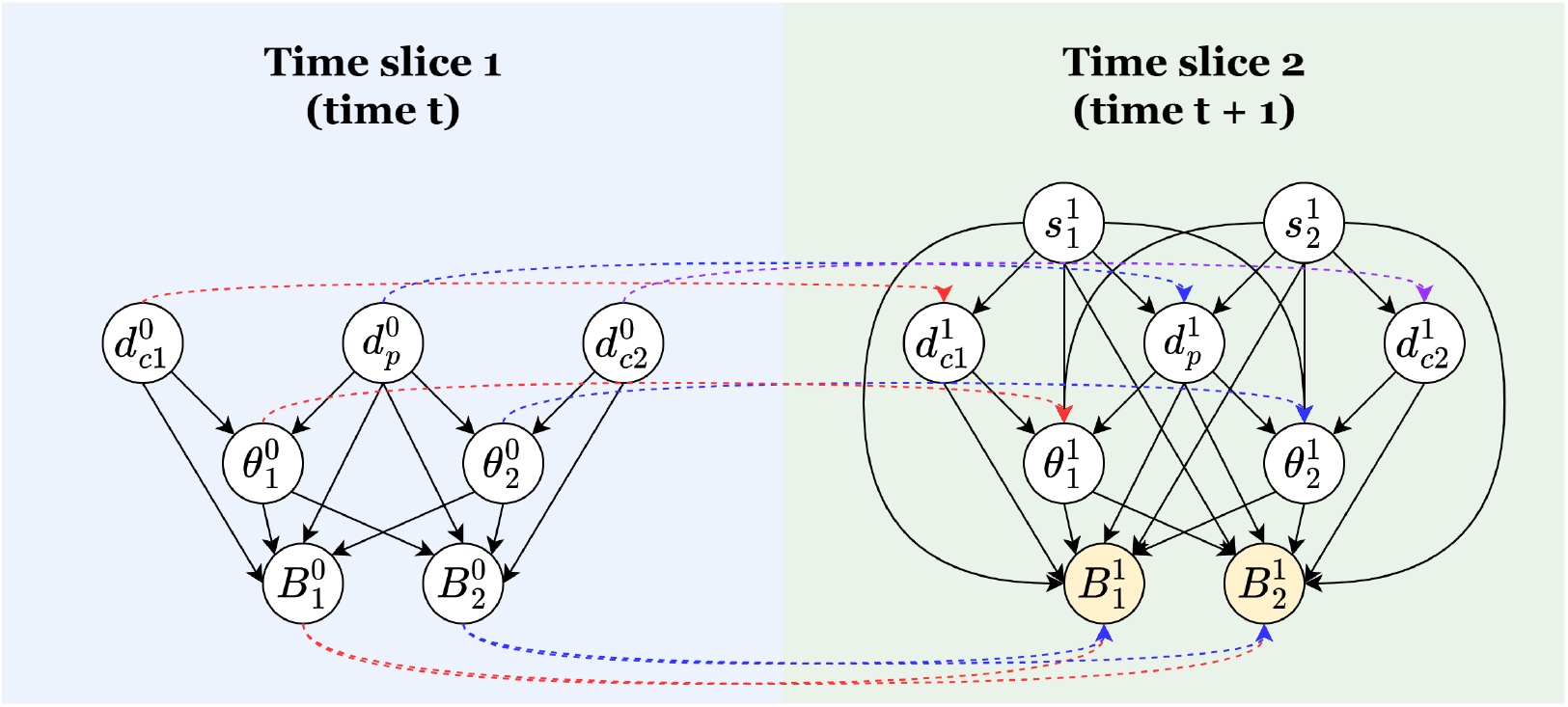
Proposed Network. Our proposed network has two time slices and upholds the Markovian assumption for all variables.

In the pruned model, for every node, we removed such children nodes that are also the descendent nodes of other children nodes. This was done in order to reduce the model complexity further by removing the unnecessary redundancy in paths for information flow. We name our unpruned network the base model.

## Experiments

### Training and Testing Procedure

We used each of the two proposed networks as a predictor by first training it and then performing queries on the target variables by supplying the rest of the variables as evidence. The value with the maximum aposteriori probability was taken as the predicted value.

The learning dataset was split into training, validation, and testing datasets with a 60%-20%-20% split. The training data (about 32.9 million rows) was used to train the model and the validation data (about 11 million rows) was used to check the performance of the hyperparameters.

On the testing data, we predict the behavior of both the flies simultaneously at the current time step (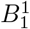 and 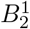) given all other variables of the two time steps, including fly behaviors of the previous time step, as shown in equation 3.

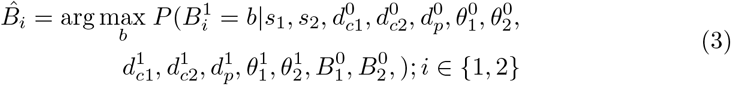

### Evaluation of the efficiency of models

The base model and the pruned model were evaluated using the following standard metrics:

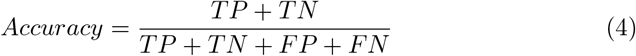

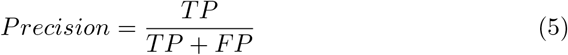

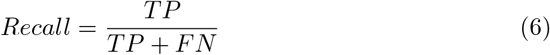

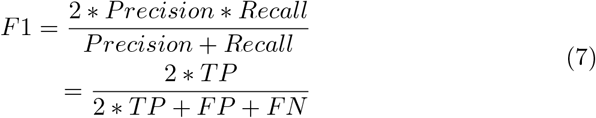

Here, TP, TN, FP, and FN are the numbers of true positives, true negatives, false positives, and false negatives respectively in the predictions.

Since this is a multi-class problem, for precision, recall, and F-1 score, we use the ‘macro’ version that calculates the metrics for each class and then takes an unweighted mean.

We evaluated these metrics for the following:

- Fly 1 Behavior
- Fly 2 Behavior
- Fly 1 and Fly 2 Behavior combined as just Fly Behavior

## Results and Discussion

The base model and the pruned model both show remarkable performance as can be observed in Table 4. Both models are able to predict the next time step behavior with good accuracy, precision, and recall. This shows that the models can correctly predict almost all instances of any behavior and of all predictions of any particular behavior, almost all are correct. Moreover, both models are able to predict almost all the instances clearly as belonging to some particular behavior or not.

**Table 4.**
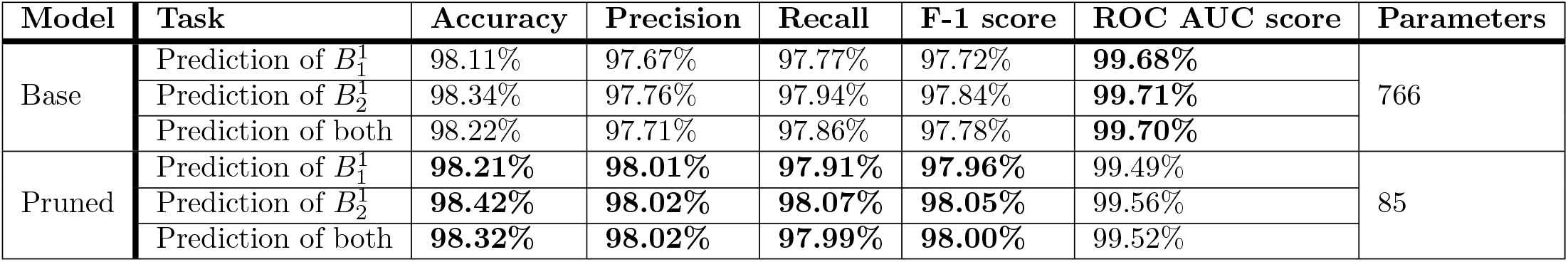
Model performance.

| Model | Task | Accuracy | Precision | Recall | F-1 score | ROC AUC score | Parameters |
| --- | --- | --- | --- | --- | --- | --- | --- |
| Base | Prediction of $B_1^1$ | 98.11% | 97.67% | 97.77% | 97.72% | **99.68%** | 766 |
| | Prediction of $B_2^1$ | 98.34% | 97.76% | 97.94% | 97.84% | **99.71%** | |
|  | Prediction of both | 98.22% | 97.71% | 97.86% | 97.78% | **99.70%** |  |
| Pruned | Prediction of $B_1^1$ | **98.21%** | **98.01%** | **97.91%** | **97.96%** | 99.49% | 85 |
| | Prediction of $B_2^1$ | **98.42%** | **98.02%** | **98.07%** | **98.05%** | 99.56% | |
|  | Prediction of both | **98.32%** | **98.02%** | **97.99%** | **98.00%** | 99.52% |  |

It can be seen that the pruned model marginally outperforms the base model. However, it is noteworthy that the models show similar performance for the prediction of the next-time behavior of either fly. Overall, the extent of error in prediction is minimal, as can be seen from the confusion matrix for the prediction task for each model in Fig 7.

**Fig 7.**
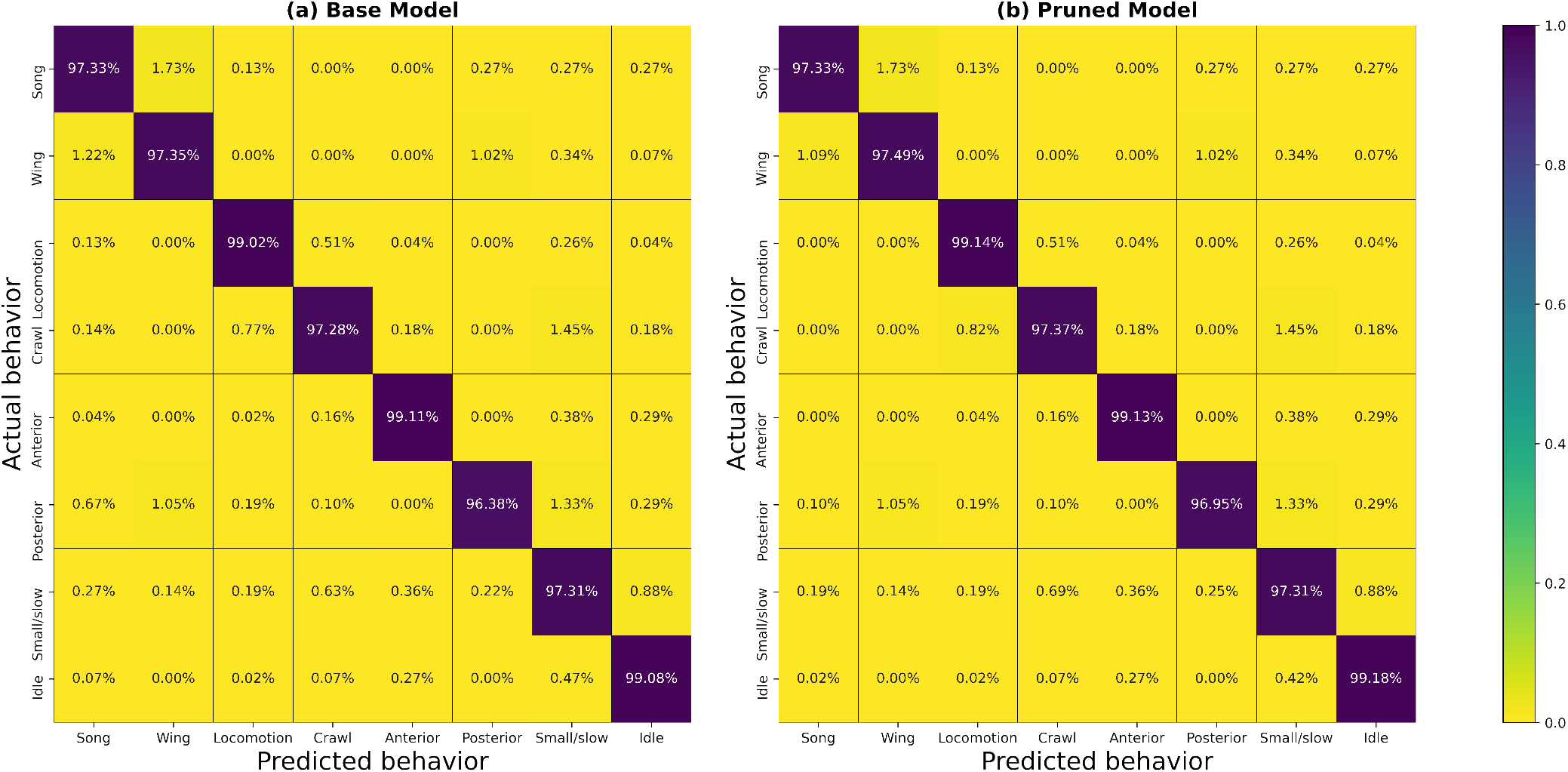
Confusion matrices. The confusion matrices show good prediction performance with very less erroneous predictions. The high values on the diagonal indicate that for the next time step, the behavior predicted by the models was almost always the same as the actual behavior. All the non-diagonal values representing misdetection are almost negligible. The models work comparably well for all behaviors considered.

In this work, for every entry of testing data, a query was performed on the model in use for the behavior given all other observed variables as evidence. This gives the probability for each behavior, of which the prediction is the behavior that has the highest probability. This approach does not use any predetermined threshold for probabilities to determine the predicted behavior. To evaluate the prediction efficiency of the models across all thresholds, Receiver Operating Characteristic (ROC) curves were plotted as seen in Fig 8 and their Area Under the Curve (AUC) scores were examined.

**Fig 8.**
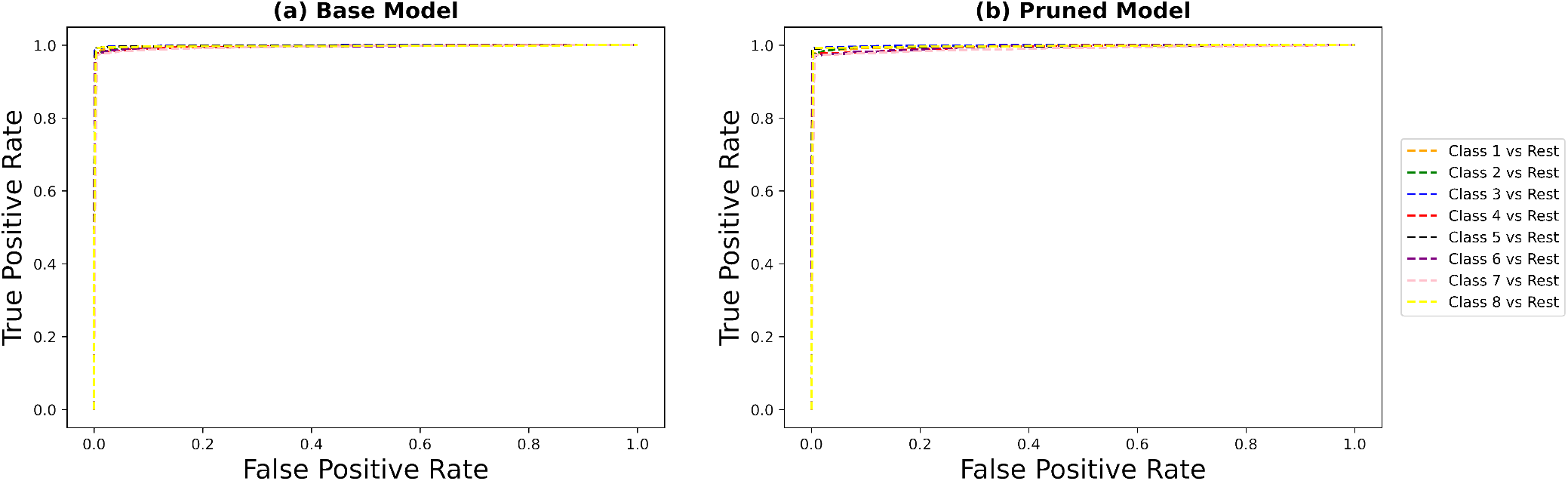
Receiver Operating Characteristics Curves. ROC curves are created by plotting the True Positive Rate (Sensitivity) versus the False Positive Rate (1-Specificity) for binary class prediction at every threshold from zero to one. The area under the curve helps evaluate the discriminative ability of any model for the class. To adapt this to our multi-class prediction task, we treat each instance of prediction as a one-versus-all prediction of each class and plot one curve for each class. For either model, the curve formed is almost perfect for all classes.

The ROC curves are near perfect and the AUC scores are close to unity. This highlights the discriminative power of the models. They are able to make every prediction with high conviction. In other words, as a result of every query, the predicted behavior has a very high probability compared to other behaviors. The pruned model has a better AUC score than the base model, which shows that the base model is a better discriminator.

### Dynamic Bayesian Network based approach as a tool for modeling

The proposed approach employing Dynamic Bayesian Networks proves to be a valuable alternative to standard approaches since it explains the domain data lucidly in a graphical yet quantifiable method. Despite such prediction efficiency, the base model is lightweight and utilizes only 766 parameters, which is a tiny fraction of what their heavier counterparts, neural networks, would require. Moreover, the proposed model is essentially learned from the data of the domain and provides a meaningful representation, unlike black-box models, without any compromise on performance. The pruned model has an 88.9% reduction in the number of parameters from the base model yet performs better than the base model. Such performance and efficiency highlight the strength of the proposed network to model the exhibited domain in an explanatory manner. In general, models with fewer parameters (simple) are preferred over models with larger parameters (complex) if the simpler model better fits the data at hand (Occam’s razor).

Understanding the nuances of animal behavior is challenging. Any behavioral outcome is a likely result of several interacting factors. Rather than addressing behavior in parts, a comprehensive approach to predictive modeling could be helpful. Even then, interpretations of behavioral analyses seldom address finer spatiotemporal patterns. To gain a more holistic view, we used dynamic Bayesian modeling which could take into account spatiotemporal complexity and still predict behavior in paired interactions well. To our knowledge, this issue has not been addressed in previous studies attempting modeling of behavior or social interactions.

Even though a molecular toolkit for *Drosophila* is available, several Drosophilid species’ social interaction between two individuals is biased by genetic variation and context. Thus, there might not be repeatability in the patterns observed in a given set of behavior. Slightly better predictability could be aimed at the mating ritual compared to male-male or female-female interactions. Yet, mating rituals could also vary across species; in terms of the approach of the male towards the female (alternative to approaching from the back). Our study could be used as a template to model known behavioral observations and corroborate existing results on social behavior. Significant literature details behavior. It is quintessential to look into it through a predictive lens and that too an unbiased one.

## Conclusion

We propose a novel approach to model and predict the behavior of a pair of *Drosophila melanogaster*. The models are symmetrical to allow random assignment of fly numbers, overcome the acyclicity obstacle by being modeled as a 2-time-slice Dynamic Bayesian Network, and very lightweight as they require a very small number of parameters. It is worth noting that they can be easily modified for different tasks and scenarios by the same approach. Including more time slices in the proposed models allows us to predict the behavior after a multiple number of time steps. The same method can be used to form models for multiple flies with the appropriate dataset. Our models improve significantly over earlier attempts by being capable of handling complex spatiotemporal patterns of behavior, modeling the paired social contexts, and providing a quantified explanation of the relationship between various variables considered. Moreover, our models show impressive performance in the prediction of fly behavior at the next time step.

